# Electronic Actuation of Surface-Immobilized, pH-Responsive DNA Nanoswitches

**DOI:** 10.1101/2025.07.17.665395

**Authors:** Francisca D’Rozario, Callum D. Silver, Katherine E. Dunn, Steven D. Quinn, Andy M. Tyrrell, Christoph Wälti, Steven Johnson

## Abstract

Dynamic DNA machines exploit the specificity of base pairing and/or sensitivity to the local environment to control the reversible switching of DNA constructs between conformational states. One such example are pH-sensitive DNA nanoswitches that can be actuated by proton-mediated Hoogsteen interactions within a DNA triplex domain. To date, studies of pH-sensitive DNA nanoswitches have largely focused on DNA machines that are freely diffusing in the solution phase. For many applications, it is advantageous to integrate these dynamic DNA machines with solid-state devices, requiring immobilization on surfaces. Here, we explore the switching of a pH-sensitive DNA triplex immobilized on a surface as a dense, 2-dimensional DNA monolayer. DNA nanoswitches were assembled onto surfaces via thiol chemistry and pH-controlled conformational switching of the constructs examined using quartz crystal microbalance with dissipation monitoring (QCM-D). These QCM-D experiments indicate that despite the high density of DNA within the monolayer (10^12^ molecules/cm^2^), pH-switching between open and closed states is retained following immobilization. Moreover, conformational switching of DNA constructs within the monolayer remains highly reversible and repeatable, with negligible reduction in switching efficiency observed over 20 switching cycles. DNA switching experiments were also performed in the solution phase using single-molecular Förster resonance energy transfer (smFRET) and circular dichroism (CD) techniques to confirm their pH responsitivity. Finally, we demonstrate electrically driven, localised, and addressable switching of the DNA triplex by employing electrochemical reduction and oxidation of water at an electrode surface, further demonstrating the potential of the technology for surface-immobilized dynamic DNA machines. This study not only provides insight into the actuation of DNA machines on-surface but also supports the development of new technologies such as hybrid electronic-DNA technologies able to store and process information using both molecular and electronic inputs.

## INTRODUCTION

DNA triplexes comprising three strands of oligonucleotides are a well-studied example of the variety of non-canonical structures that DNA molecules can adopt [1]. DNA triplexes consist of a double stranded domain stabilised by Watson-Crick and a third oligonucleotide sequence that binds to the major groove of the double stranded domain via Hoogsteen hydrogen bonding interactions **(Figure 1)**. The base-pairing schemes in DNA triple helices are *T*AT, *A*AT, *C*GC and *G*GC (third-strand bases in italics). Importantly, Hoogsteen interactions are strongly dependent on local pH. Specifically, *C*GC triplets are favoured at acidic pHs due to protonation of the N_3_ of cytosine on the third strand (pKa of protonated cytosines in triplex structure is ≈ 6.5). *T*AT triplets are stable at neutral pH but become unfavourable at basic pH due to deprotonation of thymine (pKa ≈ 10). This leads to a dependency between DNA triplex stability and local pH, where the pKa of the triplex depends on the number and ratio of TAT and CGC triplets [2,3] and that can be exploited to create DNA constructs that behave as switchable nanomachines, or nanoswitches for use as logic gates for molecular computation [4], dynamic building blocks of DNA origami [5], components of drug-delivery vehicles [6], and synthetic machines to regulate gene expression [7].

**Figure 1.**
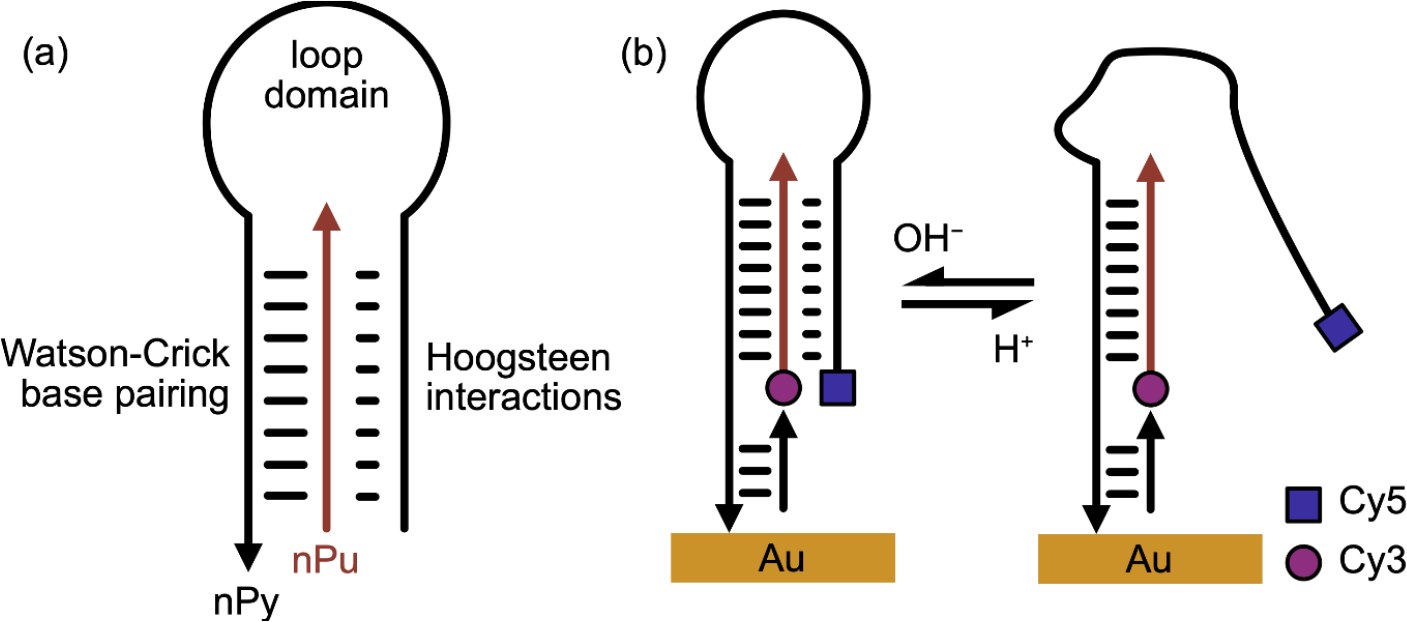
Surface-immobilised DNA triplex nanoswitch construct (arrowhead indicates 3’end). (a) A simplified intermolecular DNA triplex switch constructed from a polypurine strand (pPu) and a polypyrimidine (pPy) strand, shown here in the closed state (at pH < 7) where the conformation is stabilised by both Watson-Crick base pairing and Hoogsteen interactions. (b) (left) The nanoswitch in the closed state, in an acidic environment. Immobilization is achieved by the inclusion of a thiol modifier at the 3’-end of the polypyrimidine oligonucleotide. A short oligonucleotide complementary to the extension at the 3’-end of the polypyrimidine strand was included to increase rigidity. Cy3 (donor) and Cy5 (acceptor) fluorophores were used to label the DNA constructs for single molecule FRET experiments. (right) The DNA switch in an open state at basic pH.

pH-dependent switching of freely diffusing DNA triplexes has largely been studied by changing the solution pH by titration [8]. It has also been shown that the state of pH-dependent DNA nanoswitches can be changed from open to closed conformations using electrochemical reactions that lead to a reversible change in bulk solution pH. For instance, Minero *et al*. used voltage-biased gold microelectrodes to cycle between pH 4 and 8 using quinhydrone redox systems [9]. With their approach, pH changes were used to control the conformation of a DNA in solution from duplex to triplex, which acted as a catalyst for a disulphide ligation reaction. Yang *et al* presented a microfabricated chip that was used to drive electrolysis of water to reversibly switch the pH between 5 and 8 on a timescale of 20 seconds per step and induce transitions in a DNA nanoswitch based on an i-motif in-solution[10].

These examples of electrically-actuated pH switching of dynamic DNA machines demonstrate the enhancement in speed, convenience and programmability that can be achieved compared to traditional titration approaches. However, they still rely on delocalized ensembles of DNA switches that are contained in and able to diffuse freely through solution. There is currently increasing interest in the operation of dynamic DNA machines on surfaces where surface-immobilization can be used to control the spatial arrangement of DNA molecules on the surface, with high spatial resolution [11]. A range of computational machines have been demonstrated that exploit this spatial control of surface-immobilised machines to either promote or supress interactions between DNA machines [12-15].

While approaches to immobilise synthetic DNA oligonucleotides onto surfaces are well-established, molecular crowding can play an important role in regulating the behaviour of surface-immobilized molecules, which are typically more densely packed than their solution-phase counterparts. For instance, DNA hybridization is known to proceed more slowly on surfaces than in solution [16] and the kinetics of toehold strand displacement are affected by surface-immobilization [17].

Despite the interest in surface-phase dynamic DNA machines, the surface-immobilized pH-induced switching of DNA triplex nanoswitches have not yet been fully investigated. To address this, we undertook a detailed study of pH switching of a DNA triplex on-surface using quartz crystal microbalance with dissipation monitoring (QCM-D) [17]. QCM-D involves the use of acoustic waves to probe surface-immobilized molecules, providing real-time information on their mass. For example, QCM-D has been used previously to explore the dynamic changes in mass of surface immobilised DNA-machines that are fuelled by toehold strand displacement [17]. Critically, QCM-D is also uniquely sensitive to the viscoelastic properties of the immobilised molecular layer, which we exploit here to explore pH-driven conformational changes occurring within a DNA triplex nanoswitch.

The DNA nanoswitches used in this study, and shown in Figure 1, are based on the design by Idili *et al*. [18] with minor modifications to enable surface immobilisation. Single molecule Förster resonance energy transfer (smFRET) and circular dichroism were first used to quantify the switching conditions in solution before QCM-D experiments confirmed that pH-switching between open and closed states is retained following immobilization, and conformational switching within the DNA monolayer is highly reversible and repeatable. Having confirmed switching on-surface, we finally investigated surface immobilization of a DNA triplex nanoswitch local to a microelectrode array to successfully demonstrate spatially localized, electronic-actuation of a DNA nanoswitch array. We anticipate that the approach demonstrated here could ultimately inform the design of hybrid systems able to perform computation with both molecular and electronic inputs [19].

## MATERIALS AND METHODS

### Buffers

Tris-EDTA (TE) buffer (1x), pH 8, was purchased from VWR International and 99% pure NaCl purchased from Sigma Aldrich was added to it to make a 1M salt solution in which the DNA triplexes were assembled. Potassium phosphate (KP_i_) buffers of pH ranging from 6 to 8.2 were used to study the pH responsiveness of the switch. The KP_i_ buffers were prepared by adding fixed volumes of potassium phosphate monohydrate (K_2_HPO_4_) and potassium phosphate dihydrate (KH_2_PO_4_) explained in section 1 of the supplementary information (SI).

### Oligonucleotides

The DNA sequences of all oligonucleotides used in this study are presented in section 2 of the SI. DNA oligonucleotides (Polypyrimidine (pPy) ssDNA strands with a thiol-modified (Thiol Modifier C3 S-S) 3’-end and cyanine 5 labelled 5’-end (Cy5-pPy-SH), and “Linker” strands, which are strands complementary to the bases near the 3’ end of the pPy strands, were ordered from Integrated DNA Technologies (IDT). Polypurine (pPu) strands with cyanine 3 labelled 5’-end (Cy3-pPu) were ordered from Eurofins Genomics. All strands were purified by IDT by high-performance liquid chromatography (HPLC), except the linker strands which were purified by polyacrylamide gel electrophoresis (PAGE).

DNA switches were assembled by mixing a 100µM 1:1:1 ratio of Cy5-pPy-SH, Cy3-pPu and Linker strands in the prepared TE/NaCl buffer (∼pH 8) at 16°C, after annealing for 5 minutes at 90°C and cooling it to room temperature overnight. Further preparation differed for each method and is described in the individual methods sections.

### CD Measurements

Measurements were performed on a Chirascan (Applied Photophysics) in 0.2 mm cylindrical, quartz cells. Scans were performed between 250 to 310 nm at a scan rate of 1 nm/second. An average of three scans was used and a buffer blank was subtracted from the raw data. DNA nanoswitches were assembled in 10 mM phosphate buffer at a range of pH (pH 5.9, 7.2, 7.7 and 8.8) at a final concentration of 25 µM.

### Single-Molecular FRET by Confocal Volume Microscopy

The smFRET experiments were carried out using an EI-FLEX smFRET spectrometer purchased from Exciting Instruments (theory in SI, Section 4). Briefly, 70 μL droplets containing ∼20 pM DNA triplex in TE + 1M NaCl were pipetted onto 30 mm #1 cover slips (VWR) and donor and acceptor emission bursts were collected under 520 nm and 638 nm alternating laser excitation (ALEX). The excitation powers were 0.35 mW and 0.17 mW, respectively and ALEX was performed using a laser alternation period of 100 µs. Here, the 520 nm laser was on for the first 45 µs, followed by a 5 µs window where both lasers were off. The red laser was then switched on for 45 µs, followed by a second 5 µs window where both lasers were off. This cycle was repeated for the duration of the measurement. Two avalanche photodiodes collected the photon arrival times for 60 minutes, with a burst rate of approximately one burst per second. Stoichiometries and apparent FRET efficiencies were then quantified using FRETBursts. [20].

The EI-FLEX spectrometer detects the donor and acceptor emissions from single molecules as they diffuse through the confocal volume, under alternating laser excitation (ALEX). E, the apparent efficiency, is calculated as:

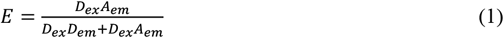

And stoichiometry, S, a measure of dye presence is given by:

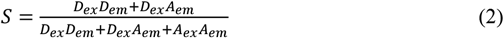

where D_ex_A_em_ is the acceptor emission under donor excitation, D_ex_D_em_ is the donor emission under door excitation and A_ex_A_em_ is the acceptor emission under acceptor excitation (Figure S3). A low E with a high S suggests donor-only molecules, whereas a low S suggests acceptor-only molecules. Intermediate S results from doubly labelled molecules.

### Quartz Crystal Microbalance with Dissipation

QCM-D experiments were conducted using a Q-sense E4 system and gold-coated sensors (QSX301, fundamental frequency 5MHz), both from Biolin Scientific. Sensors were cleaned prior to each experiment using the procedure established previously [15]. 500 nM of assembled DNA switches were used for the experiments along with 1mM MCH (mercapto hexanol) also prepared in the same buffer. 500 nM Cy5-pPy-SH strands were used as controls. The thiolated DNA construct was immobilised onto the sensor surface by flowing a DNA solution in 1M NaCl/1 × TE over the sensor surface for 15 mins at a constant flow rate of 75 μL/min using a peristaltic pump. Next, a freshly prepared solution of 6-mercaptol-1-hexanol (MCH, 97% pure from Sigma Aldrich) in 1M NaCl/1 × TE was delivered to the sensor surface at a constant flow rate 75 ul/min except. This backfilling agent was used to force the DNA switches to immobilize in an upright position and reduce non-specific binding to the gold surface [21]. The DNA construct was fully assembled prior to the experiment by incubation in solution at room temperature.

The mass density of DNA switches immobilized on the sensor surface, Δm (in ng cm^−2^), was calculated via the Sauerbrey equation (Eq. 3).

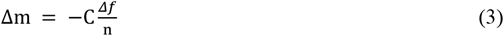

Where C is a constant equal to 17.7 ng cm^−2^ s^−1^ and n is the overtone number. Typically, measurements are made for several overtones (resonant frequencies) where the lowest overtone penetrates furthest into solution. In this paper, data is presented for the ninth overtone only.

### Electrode-based switching experiments

Electrode-based switching experiments were carried out using quartz glass disks patterned with platinum electrodes. Prior to each experiment, the disks were cleaned by submerging in piranha solution (sulfuric acid and hydrogen peroxide at a 3:1 ratio). Once clean, the immobilisation of the thiolated DNA strands onto the quartz glass was carried out as follows. The disks were then left for 16 hours in (3-Mercaptopropyl) trimethoxy silane (MPTS) (4% v/v in 2-propanol) to form a thiolated silane layer on the quartz. The thiolated quartz was then placed into a solution of 15mM Copper(II) sulphate [22] for 15 min and then rinsed in DI water and dried with nitrogen. The DNA triplex was formed in solution prior to immobilization to a final concentration of 2 μM by mixing equal parts of the Cy5-pPy-SH, Cy3-pPu and linker strands in the aforementioned TE/NaCl buffer. This solution was then incubated for 15 min at 45 °*C* to allow for the complex to form. 2 μL droplets of the formed triplex were then pipetted onto the quartz surface within the electrode structure and allowed to incubate for 90 mins in a dark, humid environment.

Following DNA triplex functionalization, the quartz electrode disk was mounted into a clamped PDMS microfluidic system (SI), which allowed liquids to be exchanged across the sensor surface. Initial confirmation of the fluorophore triplex switching was carried out using 50 mM potassium phosphate buffers at pH 6 and pH 8, the data for which is shown in SI Figure S5.

Electrically-driven switching was carried out by applying a constant current using a Keithley 2400 DC source measurement unit. The fluidic chamber was filled with 500mM NaSO4 and either +30 µA or - 30 µA was applied between the surrounding electrode and the larger counter electrode. Fluorescence was then measured using a scanning confocal microscope exciting at 633 nm and measuring at 650 nm with a 40 µs/Pixel exposure time. One frame of the image took about 1.5 seconds to complete.

## RESULTS AND DISCUSSIONS

### Behaviour of the triplex switch in solution-phase

Figure 1 illustrates the DNA nanoswitches used in this study. In contrast to the unimolecular design employed by Idili *et al*, the triplexes used in our study were assembled from two separate oligonucleotides, namely a polypurine strand (pPu) and a polypyrimidine (pPy) strand which was modified at the 3’-end to permit immobilization on surfaces. The nanoswitch was designed to adopt an ‘open’ conformation at slightly basic pH due to deprotonation of cytosines in the triplex forming domain of the nPy strand. (Figure 1a). At more acidic pH, the nPy strand folded over and bound to the double-stranded region through Hoogsteen interactions, forming a triplex, ‘closed’ state (Figure 1b).

Prior to immobilisation of the nanoswitch, pH-mediated switching of the DNA between open and closed states in solution was confirmed using single molecule Förster resonance energy transfer (smFRET) at room temperature (20°C). The two oligonucleotides, pPu and pPy, were labelled with FRET pairs Cyanine3 (Cy3) and Cyanine5 (Cy5), respectively [2], such that when folded into a triplex (closed state), the dyes would be positioned close to one another (about 1nm) [23], resulting in high FRET efficiencies (E). This can be seen in Figure 2a where at pH < 7.7, smFRET measurements revealed a dominant population of DNA nanoswitches that exhibited a FRET efficiency close to 1.0, indicative of the DNA nanoswitch in the closed conformation. In contrast, at high pH (pH > 8), the triplex domain becomes unstable, leading to opening of the DNA nanoswitches and separation of the fluorophores and a corresponding reduction in FRET efficiency (∼0.2), as shown in Figure 2d. At intermediate pH (Figure 2b), smFRET revealed the presence of two, conformationally distinct populations, one associated with the DNA nanoswitch in the closed state (with high FRET efficiency) and one associated with the open state (low FRET efficiency). The number of molecules in the closed conformation was seen to be approximately equal to the number of molecules in the open conformation when the solution pH ≈ 7.83 (Figure 2c), which we define as the solution-phase dissociation pH, pK_a_^SOL^ [24]. This differs from previous solution-phase measurements of pH-dependent DNA triplexes with 60% *T*AT content in literature, where pK_a_^SOL^ = 7.5 [18], presumably because the experiments in literature were conducted at 25°C, which being a higher temperature, might affect the switching dynamics, as shown by on-surface experiments described in the next section. A FRET stoichiometry (S) vs E plot is shown in the SI (Figure S3a) to demonstrate that the data presented has been derived only from doubly labelled DNA switches.

**Figure 2.**
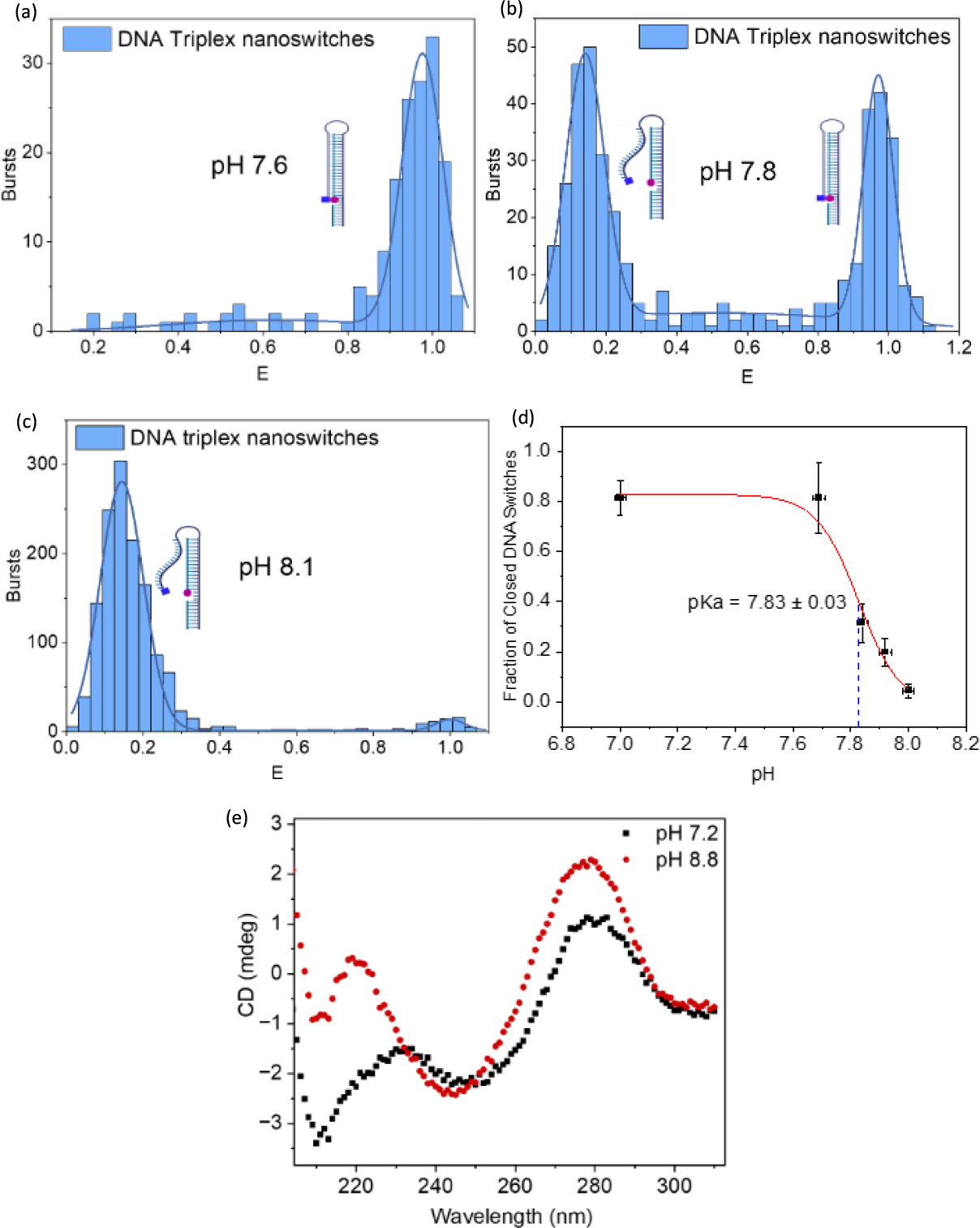
(a) Number of FRET bursts vs FRET efficiency (E) of doubly-labelled DNA triplex nanoswitches at pH 7.6 where the dominant population assumes a conformation in which the nanoswitch is closed and thus exhibited high FRET efficiency. A gaussian is fitted to the data for calculating the fraction of open and closed DNA switches. (b) Two conformational populations exist at pH 7.8 corresponding to DNA nanoswitches with high FRET efficiency (closed state) and low FRET efficiency (open state). (c) At pH 8.1, the dominant conformation of DNA nanoswitches is one in which the FRET efficiency is low, corresponding to the closed state. (d) Graph showing the decrease in population of closed switches with increasing pH, indicating the pKa as pH 7.83. The y-errors indicate the standard deviations of the gaussian fits and the x-errors denote the instrumental error of the pH-meter. (e) CD spectra at 20°C for the DNA nanoswitch in the open conformation (pH 8.8) and closed conformation (pH 7.2).

pH-mediated switching in solution was also verified by circular dichroism (CD) spectroscopy. CD spectra of unlabelled DNA nanoswitch as a function of pH are shown in Figure 2(f). At pH 8.8, the CD spectrum is characterized by large positive CD absorbance band at 276 nm, a negative band at 242 nm, and a small positive band at 221 nm. This pattern is consistent with B-form duplex DNA [25] and associated with the double stranded region of the DNA nanoswitch. As the pH is decreased, the 276 nm band reduces in magnitude and shifts to higher wavelength such that at pH 7.2, the band has shifted to 279 nm. We also observed the emergence of a negative band at 213 nm which is consistent with the existence of triple-stranded DNA at low pH values [25]. CD spectroscopy thus not only confirms pH mediated switching in solution, but also that the conformational change between closed and open states is associated with the pH-mediated assembly and disassembly of a DNA triplex. Graphs showing pH and temperature dependence are shown in the SI (Figure S1 and S2).

### Behaviour of the triplex switch on-surface

QCM-D data depicting the change of resonant frequency and dissipation with respect to time are shown in Figure 3. A change in QCM-D frequency is reflective of mass changes occurring on the sensor. Specifically, a decrease in the resonant frequency is indicative of an increase in mass, for example due to physisorption of the disulfide-modified DNA nanoswitch to the gold-coated QCM-D sensor surface, as seen in Figure 3a. From the change in frequency between the injection of DNA and saturation of the gold-coated sensor surface, the density of DNA immobilized on the gold sensors was calculated using the Sauerbrey equation (Eq. 1) and found to be of the order of 10^12^ molecules/cm^2^. This is in good agreement with published values for the density of a double-stranded DNA monolayer [17] but we note, the Sauerbrey equation assumes a uniform and rigid molecular film. After assembly of the DNA monolayer, the sensor was exposed to MCH to block exposed areas of the sensor surface.

**Figure 3.**
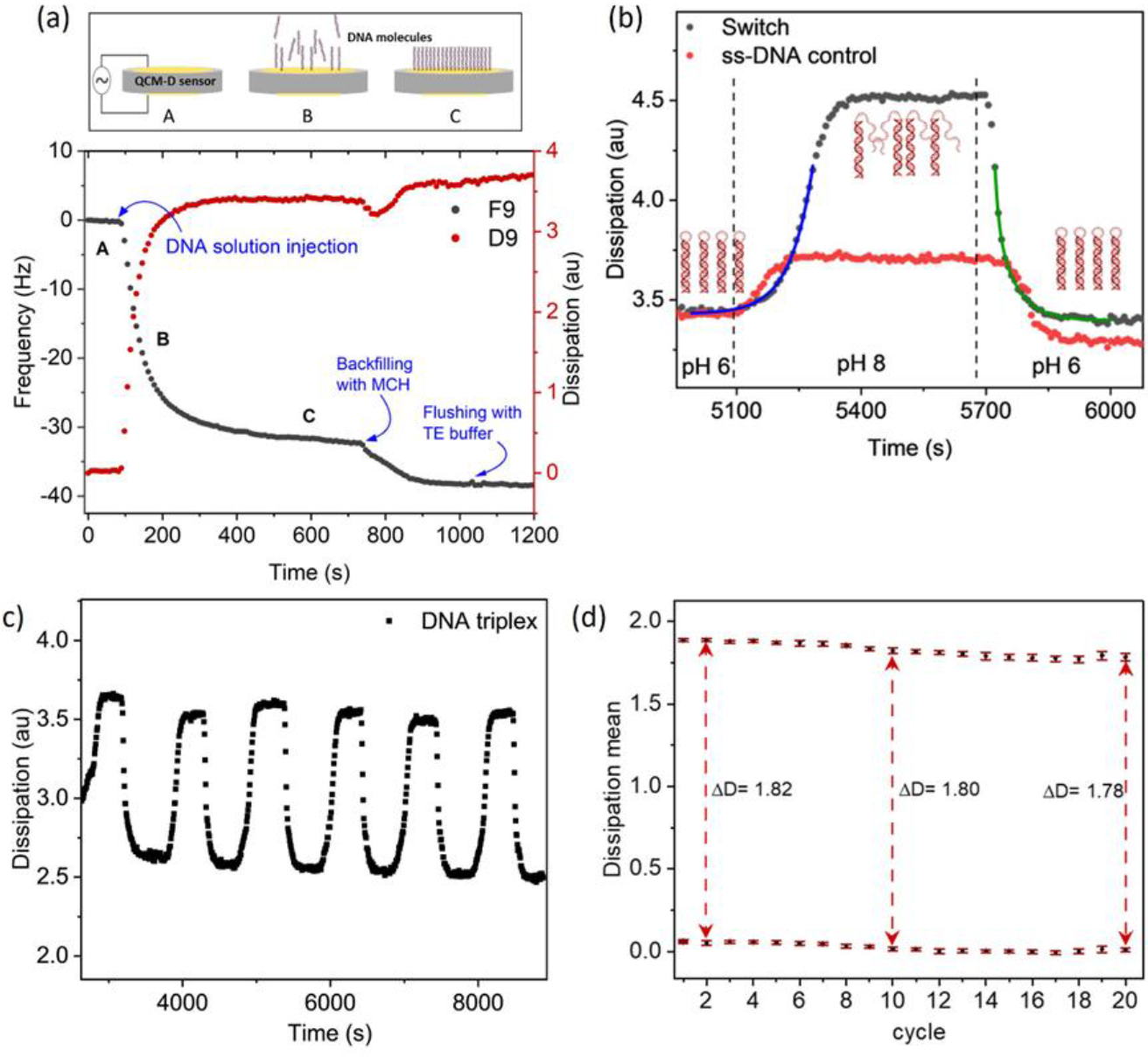
(a) QCM-D data showing changes in resonance frequency and energy dissipation during immobilisation of the disulfide-modified DNA nanoswitch on the gold-coated sensor surface. After the sensor is rinsed with TE buffer (A), the DNA nanoswitch is immobilized on the surface (B) until reaching saturation (C). The sensor is rinsed again with TE before the sensor is exposed to MCH to block exposed areas of the sensor surface. (b) The immobilized DNA nanoswitch is subsequently exposed to cycling of the solution pH by exposing to KP_i_ buffers at pH 6 followed by pH 8, before returning again to pH 6. The changes in dissipation observed at the different pH solutions indicate conformational switching between the open state at pH 8 and the closed state at pH 6. Solid curves are exponential fits to the pH-induced opening and closing of the immobilised DNA nanoswitch. Changes in dissipation for a single-stranded DNA control show the baseline shift in dissipation as a result of changes in buffer composition. (c) Switching behaviour of DNA triplexes cycling between closed states (low dissipation) at pH 6 and open states (high dissipation) at pH 8 showing repeatability and reversibility for 6 KP_i_ buffer cycles. (d) High repeatability demonstrated over 20 independent switching cycles.

The DNA functionalised sensor was subsequently exposed to repeated cycling of KP_i_ buffers of pH 6 and pH 8, which in solution was sufficient to induce conformational changes in the DNA nanoswitch from fully closed to fully open states, respectively. As shown in Figure 3b, upon changing from pH 6 to pH 8 an increase in dissipation that recovered upon returning to pH 6 was observed. In QCM-D, dissipation is a measure of the energy dissipated by the acoustic wave and is sensitive to the viscoelasticity of a surface-bound molecular layer, where an increase in dissipation is indicative of an increase in viscoelasticity. Here, the immobilized DNA switches form a more rigid, triplex conformation at pH 6 compared to the open state at pH 8 where the triplex-forming domain becomes single-stranded and thus highly flexible, dissipating more energy than when bound in a triplex. The dissipation of a QCM-D sensor functionalised with single-stranded DNA was also seen to shift when changing between pH 6 and pH 8 buffers although the magnitude of the shift was around 7 times smaller than observed for the DNA nanoswitch. This baseline shift is the result of the bulk properties of the buffer. Specifically, it has been shown that frequency [26] and dissipation [27] baselines are affected by changes in density and viscosity of the solution above the sensor, and it is known that the buffer composition, such as salt concentration, can affect both of these properties.

The triplex folding and unfolding timescales were determined by fitting exponential curves to the dissipation curves (Figure 3b). Closing of the nanoswitch was found to be best described by a second-order exponential process (with time constants of t_1_ = 11 ± 2s and t_2_ = 58 ± 5s), calculated as an average from five experiments. This agrees with thermodynamic models of DNA triplex folding that assume two rate-limiting steps: looping of the triplex-forming domain to form the first triplets, followed by zipping of the triplex and reshaping the loop region to form a stable structure [28]. In contrast, opening of the nanoswitch was found to be a single exponential process with an average time constant of 69 ± 2s, determined by the rate of dissociation of triplex-forming domain following deprotonation of cytosine in the high pH environment.

The repeatability of pH-induced DNA switching on surface was also explored by cycling the KP_i_ buffer between pH 6 and pH 8 (Figure 3c). The mean and standard deviation of the dissipation for each plateau corresponding to each KP_i_ buffer step plotted as a function of cycle number is shown in Figure 3d). The duplicate measurements agreed very closely, with the difference in dissipation between open and closed states differing by less than 2.5% across 20 cycles. The data confirms that the nanoswitch can be driven repeatedly from open to closed states without any significant deterioration for at least 20 cycles.

To evaluate the pH at which triplex dissociation occurs on-surface, pK_a_^SURF^, a sensor functionalized with DNA nanoswitches was exposed to a series of Universal Buffer solutions [29] of pH ranging from 7.0 – 9.3. Here, pK_a_^SURF^ was defined as the pH at which the observed shift in dissipation lies halfway between the dissipation < pH 7 (where it is assumed that most nanoswitches within the DNA monolayer are in the closed state) and the dissipation at pH > 8.5 (where the DNA nanoswitches are assumed to be completely open). pK_a_^SURF^ was found to be pH 8, as determined by measuring the dissipation, D, for each plateau for each of the KP_i_ buffers (Figure 4a) and calculating the pH at which 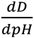 was maximum (Figure 4b) [29].

**Figure 4.**
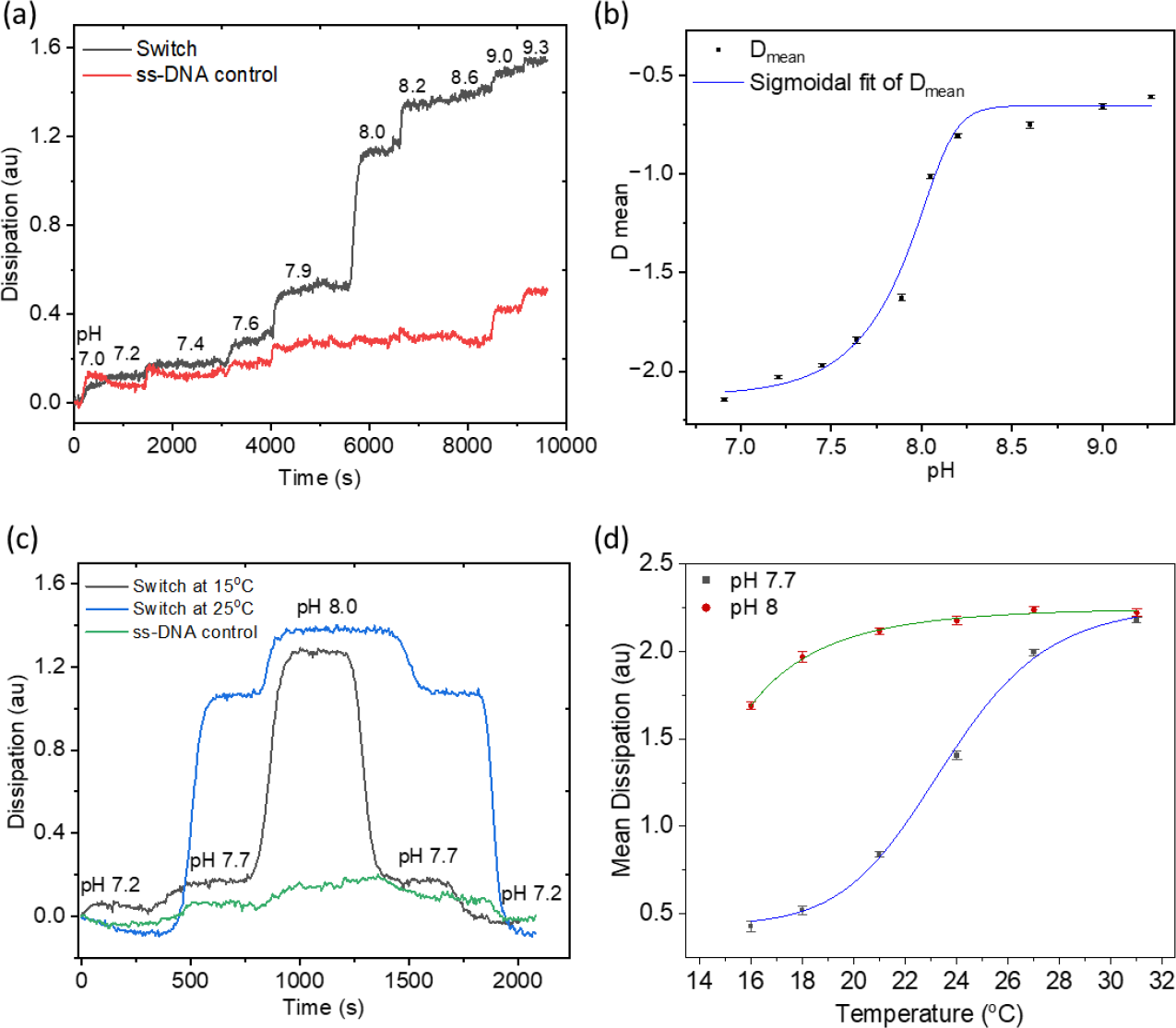
(a) Experimental QCM-D data showing the change in dissipation for a range of pH solutions between pH 7.0 – 9.3. The largest change in dissipation occurs between pH 7.9 to pH 8.0. Changes in dissipation for a single-stranded DNA control show the baseline shift in dissipation as a result of changes in KP_i_ buffer composition. (b) A sigmoidal fit to the mean of the change in dissipation for each plateau corresponding to each KP_i_ buffer step was used to identify the pH at which triplex dissociation occurs on-surface. Error bars correspond to the standard deviation of the mean of the change in dissipation for each plateau. (c) Experimental QCM-D data showing the change in dissipation as a function pH at 15°C and 25°C. (d) A sigmoidal fit of the mean dissipation values at pH 8 and 7.7 at different temperatures. The derivative of the pH 7.7 curve was used to measure the melting temperature as 23°C. Since pH 8 marks the transition pH for triplex formation at 16°C, the dissipation values saturate sooner at pH 8 than pH 7.7 with rising temperature.

**Figure 4.**
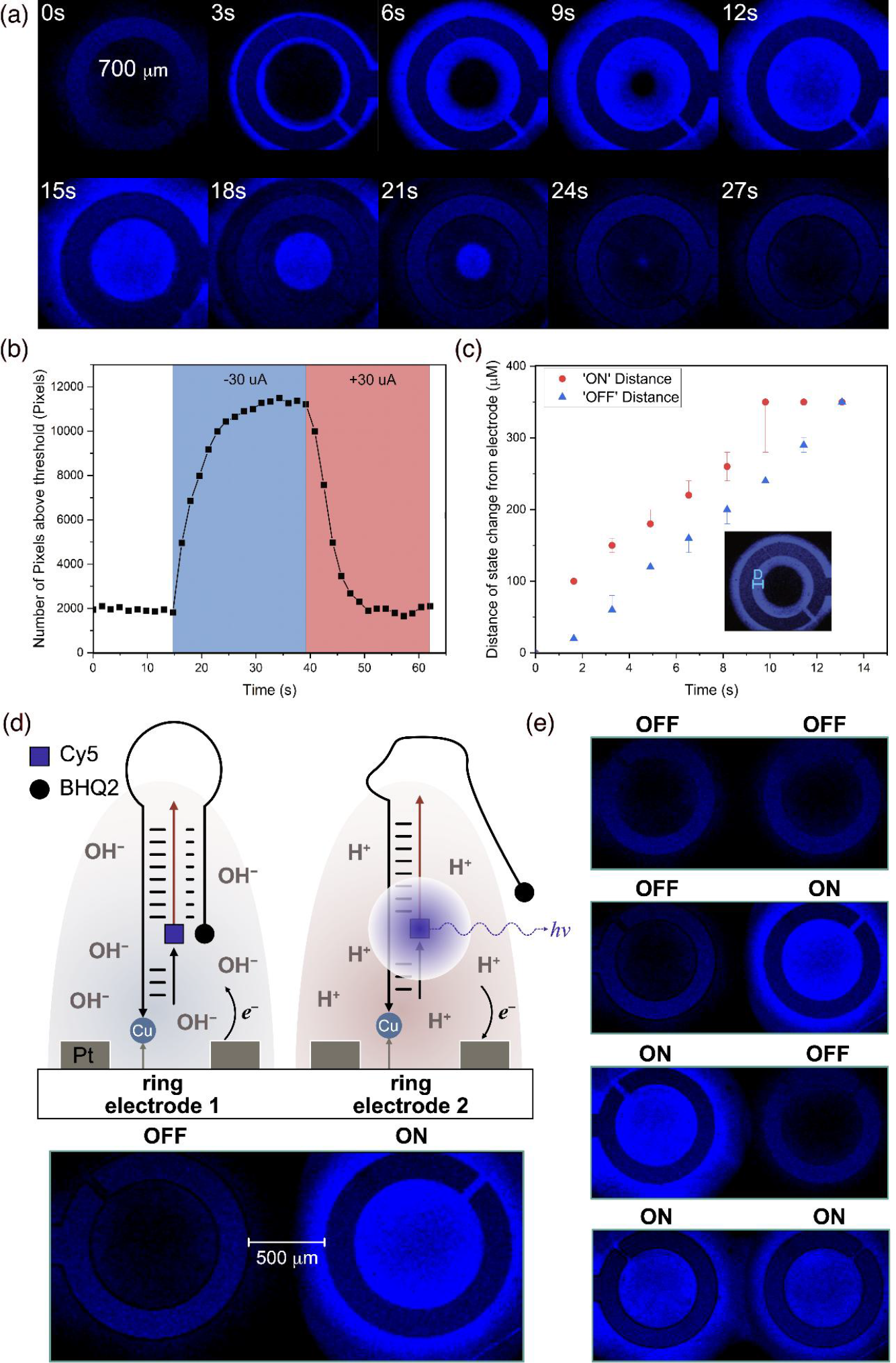
(a) Confocal microscope image demonstrating electronic actuation of a fluorescently labelled DNA nanoswitch immobilised local to a Pt ring electrode (shown schematically in panel d)). A constant current of −30 μA applied between 0 ≤ t ≤ 12 s increases the pH at the electrode surface, switching the triplex to the open conformation (or ON state) (see panel d), before being reversed to drive the anodic reaction between 15 ≤ t ≤ 27 s, to close the DNA nanoswitch. (b) Plot showing the number of ‘ON’ pixels within a single ring electrode over time for a DNA nanoswitch. ‘ON’ pixels are defined as pixels with an intensity over 110 au. (c) Speed of switching was determined by plotting the position of the ‘ON’ and ‘OFF’ pixels relative to the edge of the ring electrode (represented by D in the insert) as a function of time. (d) Schematic diagram showing immobilisation of fluorescently-labelled DNA nanoswitches local to an array of ring electrodes fabricated on a common substrate. Here, each electrode can be biased independently to locally and reversibly switch the conformation of the surface-immobilised triplex switches. e) Confocal microscope images demonstrating spatial and reversible control of surface-bound DNA triplex switching using an array of two individually addressable ring electrodes.

Finally, the thermal stability of the surface-immobilised switch at different pH was investigated by sweeping the temperature of the QCM-D sensor between 15°C to 31°C. The KP_i_ buffers were found to maintain their pH over this temperature range which is below the melting point of the duplex domain of the DNA nanoswitch (T_m_ = 72°C as simulated by NUPACK (SI Figure S3)). At pH 7.2, the DNA switch remained in the closed state over the temperature range, indicated by the low dissipation (Figure 3c). In contrast, at pH 7.7, close to pK_a_^SURF^, the dissipation was seen to increase with temperature, indicating melting of the triplex-domain and an increase in the ratio of closed to open DNA nanoswitches (Figure 4c). The mean and standard deviation of the dissipation for each plateau corresponding to each temperature step is shown in Figure 4d. From this, the melting point of the triplex at pH 7.7 was found to be around 23°C. This value provides a working range for this design of a triplex switch for applications in electronics and biosensing.

### Electronic actuation of a DNA nanoswitch

Application of a potential difference between electrodes in an aqueous solution can lead to electrolysis of the water leading to the production of H+ and OH-ions and a change in solution pH [30]. This opens up the possibility of a hybrid electronic-DNA technology in which the state of a pH-sensitive DNA construct can be regulated controllably and reversibly by electronic actuation. To explore this, we immobilised a DNA triplex switch local to an electrode within a microfluidic electrolysis cell. The conformation of the DNA triplex switch was monitored by labelling with a fluorophore (Cy 5) and quencher (BHQ-2) on the 5’ termini of the pPy and pPu respectively. In the open state, the fluorophore and quencher become separated allowing fluorescence to be observed, while in the closed state, the fluorescence is quenched, giving an ON/OFF signal dependent on the state of the nanoswitch (see Figure 4d). The DNA nanoswitch was immobilized onto a glass substrate containing individually addressable, microfabricated ring electrodes. By sourcing a constant current between a ring electrode and a Pt-wire counter electrode, the aqueous solution local to the electrode surface can be reduced or oxidised, according to the following half-reactions:

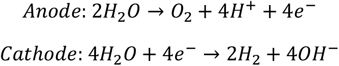

As shown, these reactions either generate additional hydrogen ions at the anode, or hydroxide ions at the cathode leading to a change in pH which is initially localised at the electrode surface. These hydrated ions can subsequently diffuse away from the surface driven by the ion concentration gradient, leading to a bulk change in pH.

Figure 4a shows the progression of fluorescence intensity over time as a constant current of −30 μA is applied to drive the cathodic reaction (between 0 ≤ t ≤ 12 s in Figure 4a) before being reversed to drive the anodic reaction (between 15 ≤ t ≤ 27 s in Figure 4a). Initially, we see a sharp increase in the fluorescence signal close to the electrode surface. Here, electrochemical production of OH^−^ ions results in a decrease in local pH that destabilises DNA triplexes immobilised close to the electrode surface, resulting in switching to the open state and an increase in fluorescence intensity. As the electrolysis reaction continues, OH^−^ ions diffuse radially away from the electrode surface, resulting in the time dependent increase in fluorescence signal away from the electrode edge (as seen between 3 ≤ t ≤ 12 s in Figure 4a). The reverse is observed when the ring electrode is subsequently biased in the opposite direction to drive acidification of the solution local to the electrode, resulting in switching of the immobilised DNA triplex into the closed state (between 15 ≤ t ≤ 27 s in Figure 4a). We note, bulk changes in solution pH by titration do not result in the distinctive time-dependent, radial change in fluorescence observed when the pH is regulated by electrolysis and ion diffusion from a localised electrode (SI Figure S5).

The fluorescent signal immediately adjacent to the electrode edge was found to reduce after repeated biasing of the electrode, potentially due to damage to the DNA and/or fluorophore by the extreme pH at the electrode surface. However, away from this region, we were able to repeatably switch the DNA between open and closed states (4 cycles shown in SI Figure S6). We note that some reduction in fluorescence intensity was observed after each switching cycle. This is likely due to fluorescence photobleaching rather than incomplete folding/unfolding of the triplex which, as shown by QCM-D experiments, demonstrated highly repeatable switching.

As shown in Figure 4b, switching of the DNA triplexes immobilised local to the electrode edge between OFF state (closed conformation) and ON state (open conformation) was observed to occur in < 2 s after application of current (corresponding to the temporal resolution of the scanning confocal microscopy measurements). Here, the ON state is defined as those pixels in which the fluorescence intensity is > 110 au). Moreover, all pixels within the 350 μm radius ring electrode are found to be in the ON state after only 15 s. Similarly, upon reversing the current, we see rapid (12 s) switching back to the closed state, corresponding to the case when all pixels are OFF (fluorescence intensity < 110 au). These rates are significantly faster than equivalent solution-based molecular switches where pH is changed by titration [9].

To further examine the reaction kinetics, we analysed the speed at which switching progresses away from the electrode edge, into the interior of the ring electrode. As shown in Figure 4c, following biasing of the electrode, the switching of pixels to the ON state and OFF state within the ring electrode is seen to occur linearly with distance from the electrode edge. By fitting these trends, we calculate that the speed at which switching progresses from the electrode edge towards the centre of the ring electrode is 22.6 ± 1.1 μ/s and 28.1 ± 2.1 μ/s for switching to the ON and OFF states, respectively (SI Figure 7), likely limited by diffusion of H^+^ and OH^−^ ions in the aqueous electrolyte.

Finally, we explore the potential of exploiting the spatial localisation (limited by diffusion of H^+^ and OH^−^ ions) of electrochemically induced pH change to demonstrate addressable switching of surface-bound DNA triplexes. As shown schematically in Figure 4d, an array of ring electrodes with a radius of 350 μm and separated by 500 μm was fabricated on a glass substrate such that each electrode could be biased independently. The nanoswitch was subsequently immobilised on the substrate surface such that the DNA monolayer covered the entire surface, including each electrode in the array. The fluorescence image of Figure 4d shows selective and independent electronic switching of DNA nanoswitches local to two adjacent ring electrodes while in Figure 4e the current is switched between the two ring electrodes in sequence to generate a 2-bit, binary sequence.

## CONCLUSIONS

We have demonstrated a DNA nanoswitch that is able to switch dynamically between open and closed conformations when immobilised on a surface. Conformational switching is regulated by the association and dissociation of a pH-sensitive DNA triplex-forming domain, as confirmed by titration in both solution and on-surface. Switching of the triplex occurs at around pH 8 when immobilised on a surface, similar that that observed for the same nanoswitch in the solution-phase, and was shown to be reversible and highly repeatable for 20 cycles of switching. By changing the pH electrochemically, we finally demonstrated electronic actuation of the surface-immobilised DNA nanoswitch. Electronically actuated switching was found to occur rapidly and switching between states was spatially localised, limited by the diffusion of ions, enabling the demonstration of electrically driven, independent and addressable switching of a DNA nanoswitch array. Further work on this system could include the development of high-density arrays containing an entire family of pH-sensitive nanoswitches, each of which is designed to respond to a different stimulus, in accordance with the work of Idili *et al* [9]. The implication of this work is the potential to integrate dynamic DNA machines with electronic systems, potentially enabling complex information processing. For example, information encoded within the loop between the duplex and the triplex-forming domain could be revealed upon opening of the switch to enable electronically-actuated searching of a key-based nucleic acid memory.

## Notes

### Competing Interest Statement

The authors have declared no competing interest.

### Funding

- Engineering and Physical Sciences Research Council: DTP studentship

## REFERENCES

[1] Kaushik, M.; Kaushik, S.; Roy, K.; Singh, A.; Mahendru, S.; Kumar, M.; Chaudhary, S.; Ahmed, S.; Kukreti, S. A Bouquet of DNA Structures: Emerging Diversity. Biochem. Biophys. Rep. 2016, 5, 388–395.

[2] Hoogsteen, K. The Structure of Crystals Containing a Hydrogen-Bonded Complex of 1-Methylthymine and 9-Methyladenine. Acta Crystallogr. 1959, 12, 822–823.

[3] Hoogsteen, K. R. The Crystal and Molecular Structure of a Hydrogen-Bonded Complex Between 1-Methylthymine and 9-Methyladenine. Acta Crystallogr. 1963, 16, 907–916.

[4] Qi, M.; et al. Reconfigurable DNA Triplex Structure for pH Responsive Logic Gates. RSC Adv. 2023, 13, 9864–9870.

[5] Sachenbacher, K.; Khoshouei, A.; Honemann, M. N.; Engelen, W.; Feigl, E.; Dietz, H. Triple-Stranded DNA as a Structural Element in DNA Origami. ACS Nano 2023, 17 (10), 9014–9024.

[6] Ijäs, H.; Hakaste, I.; Shen, B.; Kostiainen, M. A.; Linko, V. Reconfigurable DNA Origami Nanocapsule for pH-Controlled Encapsulation and Display of Cargo. ACS Nano 2019, 13 (5), 5959–5967.

[7] Knauert, M. P.; Glazer, P. M. Triplex Forming Oligonucleotides: Sequence-Specific Tools for Gene Targeting. Hum. Mol. Genet. 2001, 10 (20), 2243–2251.

[8] Idili, A.; Vallée-Bélisle, A.; Ricci, F. Programmable pH-Triggered DNA Nanoswitches. J. Am. Chem. Soc. 2014, 136, 5836–5839.

[9] Minero, G. A. S.; Wagler, P. F.; Oughli, A. A.; Mc-Caskill, J. A. Electronic pH Switching of DNA Triplex Reactions. RSC Adv. 2015, 5, 27313–27325.

[10] Yang, Y.; Liu, G.; Liu, H.; Li, D.; Fan, C.; Liu, D. An Electrochemically Actuated Reversible DNA Switch. Nano Lett. 2010, 10, 1393–1397.

[11] Germishuizen, W. A.; Walti, C.; Tosch, P.; Wirtz, R.; Pepper, M.; Davies, A. G.; Middelberg, A. P. Dielectrophoretic Manipulation of Surface-Bound DNA. IEE Proc. Nanobiotechnol. 2003, 150 (2), 54–58.

[12] Frezza, B. M.; Cockroft, S. L.; Ghadiri, M. R. Modular Multi-Level Circuits from Immobilized DNA-Based Logic Gates. J. Am. Chem. Soc. 2007, 129, 14875–14879.

[13] Smith, L. M.; et al. A Surface-Based Approach to DNA Computation. J. Comput. Biol. 1998, 5, 255–267.

[14] Liu, Q.; et al. DNA Computing on Surfaces. Nature 2000, 403, 175–179.

[15] Qian, L.; Winfree, E. Parallel and Scalable Computation and Spatial Dynamics with DNA-Based Chemical Reaction Networks on a Surface. Lect. Notes Comput. Sci. 2014, 8727, 114–131.

[16] Peterson, A. W.; Heaton, R. J.; Georgiadis, R. M. The Effect of Surface Probe Density on DNA Hybridization. Nucleic Acids Res. 2001, 29, 5163–5168.

[17] Dunn, K.; Trefzer, M.; Johnson, S.; et al. Investigating the Dynamics of Surface-Immobilized DNA Nanomachines. Sci. Rep. 2016, 6, 29581.

[18] Idili, A.; Vallée-Bélisle, A.; Ricci, F. Programmable pH-Triggered DNA Nanoswitches. J. Am. Chem. Soc. 2014, 136, 5836–5839.

[19] Dunn, K.E.; Trefzer, M.A.; Johnson, S.; Tyrrell, A.M. Towards a Bioelectronic Computer: A Theoretical Study of a Multi-Layer Biomolecular Computing System That Can Process Electronic Inputs. Int. J. Mol. Sci. 2018, 19, 2620.

[20] Ingargiola, A., Lerner, E., Chung, S., Weiss, S. & Michalet, X. FRETBursts: an open source toolkit for analysis of freely-diffusing single-molecule FRET. PLoS ONE 11, e0160716 (2016)

[21] Oberhaus, F.V.; Frense, D.; Beckmann, D. Immobilization Techniques for Aptamers on Gold Electrodes for the Electrochemical Detection of Proteins: A Review. Biosensors 2020, 10, 45.

[22] White, S. J.; Johnson, S. D.; Sellick, M. A.; Bronowska, A.; Stockley, P. G.; Wälti, C. The Influence of Two-Dimensional Organization on Peptide Conformation. Angew. Chem. Int. Ed. 2015, 54, 974–978.

[23] Hodges, C. S.; Harbottle, D.; Biggs, S. Investigating Adsorbing Viscoelastic Fluids Using the Quartz Crystal Microbalance. ACS Omega 2020, 5, 22081–22090.

[24] Idili, A.; Ricci, F. Design and Characterization of pH-Triggered DNA Nanoswitches and Nanodevices Based on DNA Triplex Structures. Methods Mol. Biol. 2018, 1811, 79–100.

[25] Plum, G. E.; Breslauer, K. J. Thermodynamics of an Intramolecular DNA Triple Helix: A Calorimetric and Spectroscopic Study of the pH and Salt Dependence of Thermally Induced Structural Transitions. J. Mol. Biol. 1995, 248 (3), 679–695.

[26] Kanazawa, K. K.; Gordon, J. G., II. The Oscillation Frequency of a Quartz Resonator in Contact with a Liquid. Anal. Chim. Acta 1985, 175, 99–105.

[27] Rodahl, M.; Höök, F.; Krozer, A.; Brzezinski, P.; Kasemo, B. Quartz Crystal Microbalance Setup for Frequency and Q-Factor Measurements in Gaseous and Liquid Environments. Rev. Sci. Instrum. 1995, 66, 3924–3930.

[28] Östling, S. and Virtama, P., A Modified Preparation of the Universal Buffer Described by Teorell and Stenhagen. Acta Physiologica Scandinavica, 1946, 11: 289-293.

[29] Davis, T. M.; McFail-Isom, L.; Keane, E.; Williams, L. D. Melting of a DNA Hairpin Without Hyperchromism. Biochemistry 1998, 37 (19), 6975–6978.

[30] Haynes, W.M.; CRC Handbook of Chemistry and Physics, 93rd Edition. 2012; Taylor & Francis: Boca Raton, FL, ISBN 9781439880494.

