## Supplementary Information for "Electronic Actuation of Surface-Immobilized, pH-Responsive DNA Nanoswitches"

### 1. Preparation of 0.1 M potassium phosphate ($KP_i$) buffer

Two solutions were prepared as follows:

Solution A: 27.2g $KH_2PO_4$ per litre (0.2 M)

Solution B: 34.8g $K_2HPO_4$ per litre (0.2 M)

The abovementioned solutions were mixed according to the following table and diluted to 200 mL with MilliQ water:

**Table S1**

| pH | Solution A (mL) | Solution B (mL) |
|---|---|---|
| 5.7 | 93.5 | 6.5 |
| 6.0 | 87.7 | 12.3 |
| 7.0 | 39.0 | 61.0 |
| 8.0 | 5.3 | 94.7 |

The solutions were diluted further with MQ water to 50mM solutions used in the experiments.

### 2. DNA sequences

pPy: 5'- TTC CTT CTT CTC TTC TCT CTC TTT GTC TCT CTT CTC TTC TTC CTT ACA CGC ATA CAC CCA T -3'

pPu: 5'- AAG GAA GAA GAG AAG AGA GA-3'

Linker: 5'- ATG GGT GTA TGC G-3'

### 3. In-solution circular dichroism

A CD vs wavelength graph plotted at different temperatures and at a constant pH of 7.2 (**Figure S1a**) shows a negative band at 213 nm at 15°C which is consistent with the existence of triple-stranded DNA, indicating a closed DNA triplex switch. As the temperature increases, this negative band at 213 nm reduces in magnitude, indicating dissociation of the triplex domain. At 75°C, this band is no longer present and the CD spectrum is instead characterized by large positive CD absorbance band at 276 nm, a negative band at 242 nm, and a small positive band at 221 nm. This pattern is consistent with B-form duplex DNA and associated with the double stranded region of the DNA nanoswitch

The melting point of the triplex also depends on the pH of the solution with the melting temperature of the triplex increasing with a decrease in the solution pH. This is seen in **Figure S1b**, where the effect of different pH environments is shown at a constant temperature of 45°C. At pH 5.9 the DNA triplexes are observed to be in the closed state indicated by the negative band at 213 nm. The magnitude of this band decreases with increasing pH, indicative of a loss of the triplex domain until above pH 7.8 when the 213 band is not longer observed and the CD spectra is instead characteristic of B-form duplex DNA and associated with the double stranded region of the DNA nanoswitch.

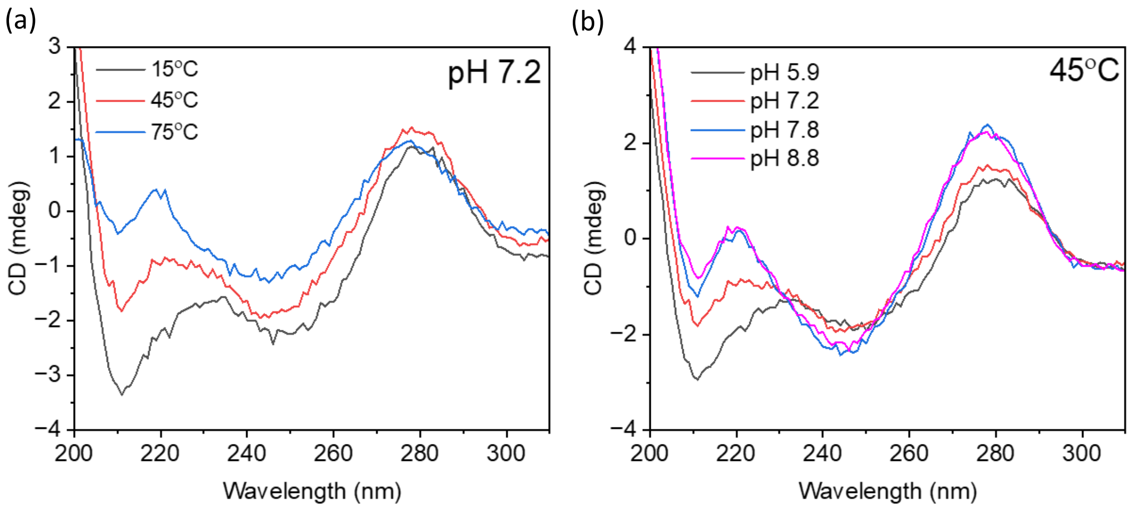


Fig S1. CD plots of DNA triplex vs wavelength: (a) At a constant pH of 7.2, DNA triplex is closed at 15°C as indicated by absorption (negative band) at the wavelength of 213 nm. The concentration of triplexes in open state increases with increasing temperature, as indicated by reduced optical absorption at 231 nm. (b) At a constant temperature of 45°C, the DNA population is predominatly in the 'closed', triplex state at pH 5.9, shifting to the 'open', duplex state with increasing pH.

The melting point of triplexes in different pH solutions was found by plotting the magnitude of the negative band at 213 nm against temperature and calculating the mid-point of the transition between high and low CD values for each curve. The melting temperatures were found to be 67°C, 46 °C, 28°C and

<10°C for pH 5.9, 7.2, 7.8 and 8.8, respectively (**Figure S2**).

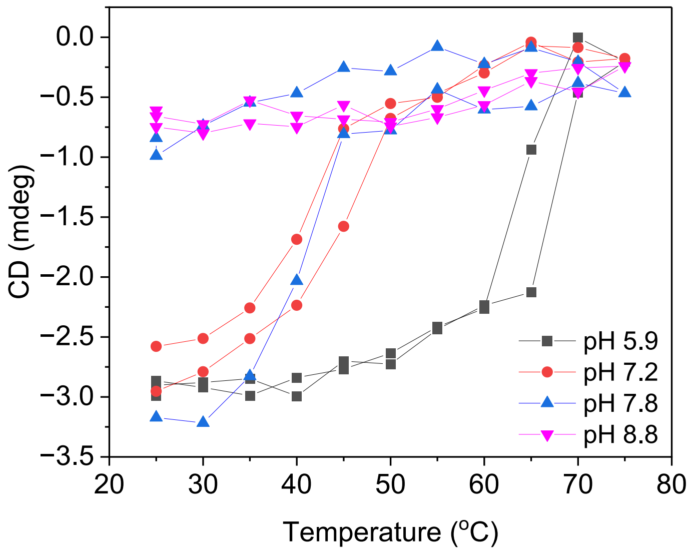


Figure S2. CD peak values at 213nm for triplexes in different pH solutions plotted against temperature.

### 4. Single Molecular smFRET

A plot of the stoichiometry (S) vs. FRET efficiency (E), centred around 0.5 depicts FRET bursts obtained by fully assembled DNA triplex molecules. Donor-only molecules show a low E and a high S. Acceptor-only molecules result in a low S, and doubly labelled molecules are depicted by intermediate S.

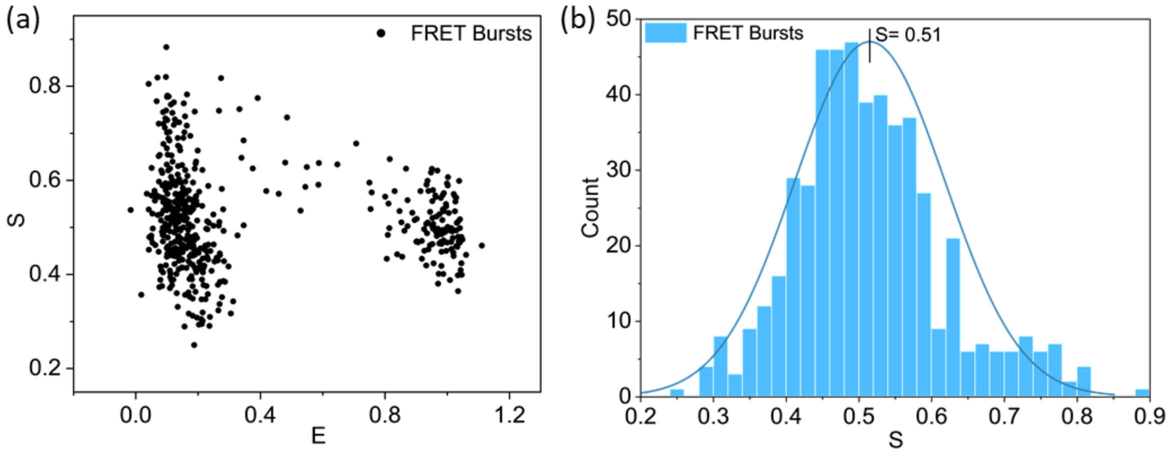


Figure S3. (a) Stoichiometry vs efficiency plot for FRET bursts of DNA switches at pH 7.92. (b) A histogram showing the distribution of the FRET stoichiometry centred at 0.51.

### 5. Melting point of the DNA switch in the "open" state

The melting point of the DNA complex containing all three strands specified in section 2 was analyzed using NUPACK, for 500 nM solution of the fully assembled DNA triplex switch. The results are shown in **Figure S4** along with the derivative which gave the peak position at 72°C, indicating the melting temperature for the DNA switch in the open state. NB. NUPACK cannot analyze triplex structures.

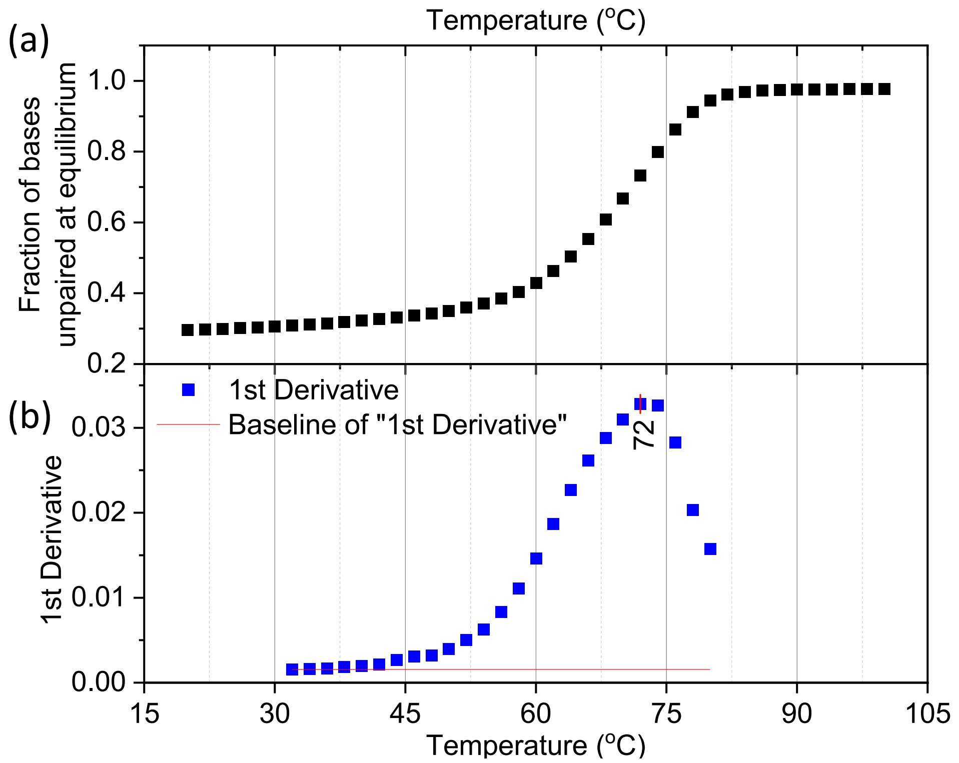


Figure S4. (a) The fraction of unpaired bases in the DNA complex plotted against temperature. (b) The derivative of the plot above, indicating the melting point at 72°C.

### 6. Electrically-controlled switching

Initial confirmation of the fluorophore triplex switching was carried out using 50 mM potassium phosphate (KPi) buffers at pH 6 and pH 8.5. By exposing the surface-bound triplexes to these pH buffers, the fluorescence was confirmed to be quenched at pH 6 and fluoresce at pH 8.5 (**Figure S5**). It was noted that the switching rate was, expectedly, faster than that of the electrode-controlled switching, showing no sign of the time-dependent, radial change shown in figure 4 (a).

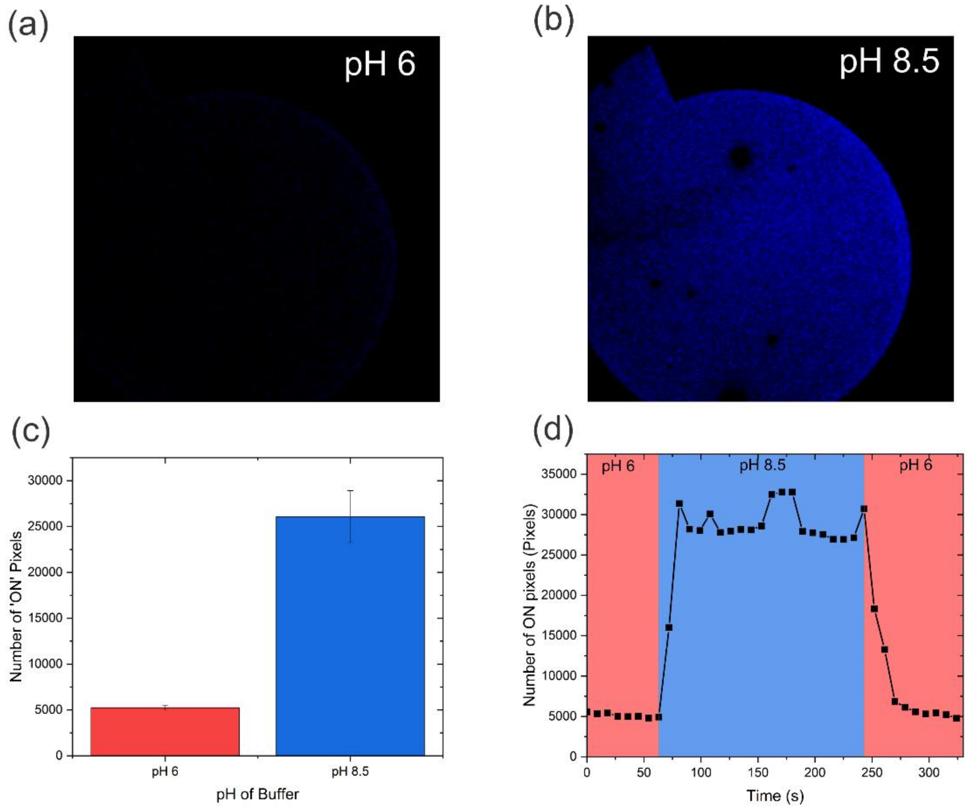


Figure S5. (a) Confocal microscope image of the fluorescently-labelled DNA-nanoswitch coated surface in pH 6 50 mM KPi buffer. (b) Confocal microscope image of the fluorescently-labelled DNA-nanoswitch coated surface in pH 8.5 50 mM KPi buffer. (c) Plot showing the average number of 'ON' pixels for each of the pH 6 and pH 8.5 KPi buffers. 'ON' pixels are defined as pixels with an intensity over 110 au. The error is reported as SD over 20 images. (d) Plot showing the number of 'ON' pixels as the pH on the surface was changed between pH 6 and pH 8.5 using KPi buffers.

The DNA switch was able to repeatably switch between open and closed states. **Figure S6** demonstrates the repeatability of DNA switching when cycling the electrode-counter current between 30 μA and -30 μA. We do, however, note that some reduction in fluorescence intensity and increase in switching time was observed after each switching cycle. We believe this is a factor of the un-controlled switching mechanism or photobleaching, rather than an inherent limitation of the DNA switch, as QCM-D experiments demonstrated highly repeatable switching.

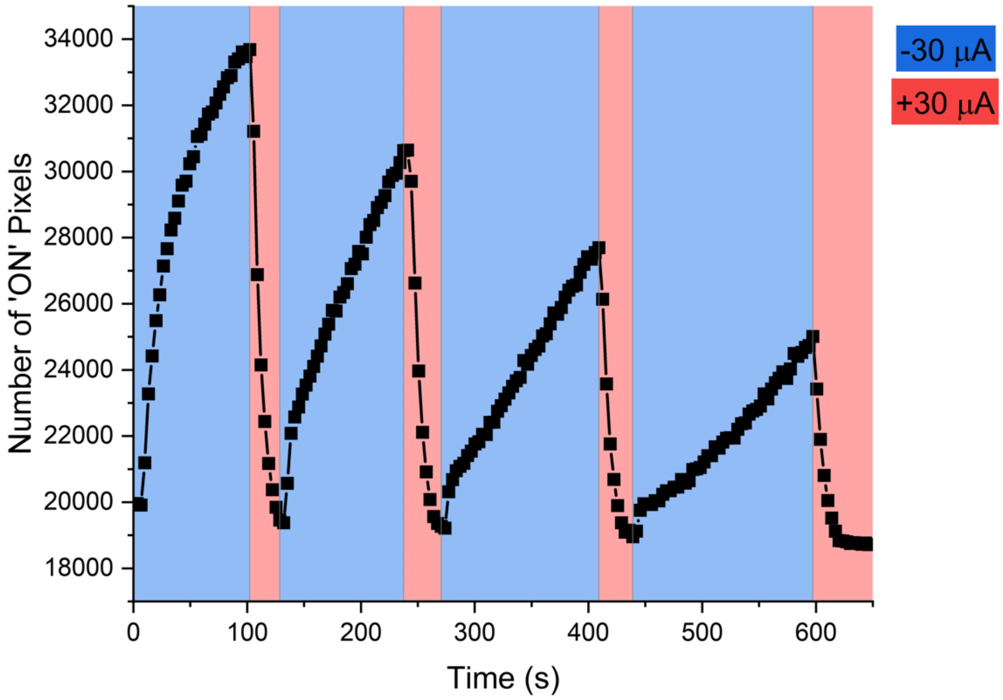


Figure S6. Plot showing the number of ‘ON’ pixels within a single ring electrode over 4 cycles of switching the DNA nanoswitch. ‘ON’ pixels are defined here as pixels with an intensity over 110 au.

To determine the speed at which switching progresses from the electrode edge, a straight line was fitted to the positions where the state of the switch changes as a function of time, giving 22.6 ± 1.1 μ/s and 28.1 ± 2.1 μ/s for switching between the ON and OFF states. Due to the relatively slow measurement speed of c.a. 2 s per image, only a portion of the data from t = 1.6 s to t = 8.2 s was used to fit the straight line – avoiding any points containing fully ON or OFF electrodes.

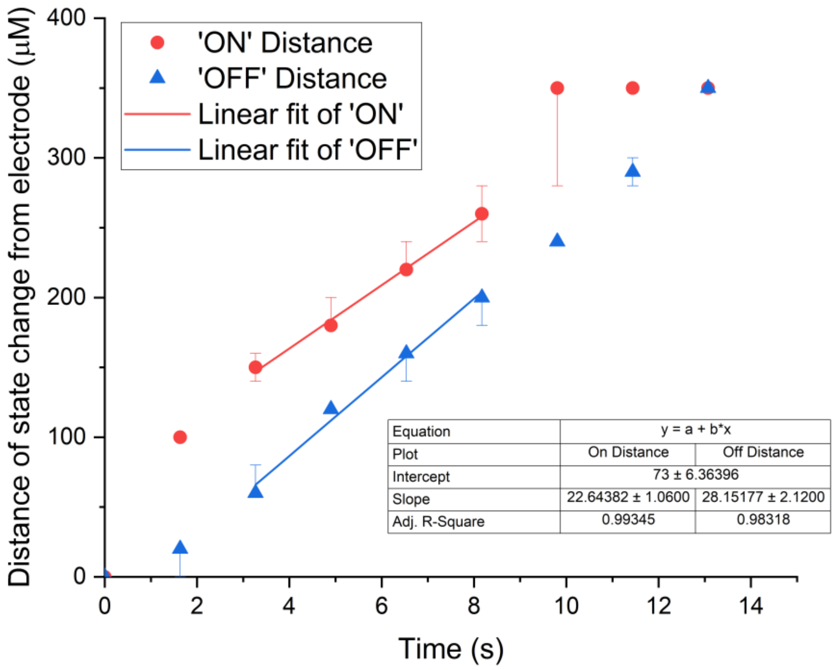


Figure S7. Speed of switching, determined by plotting the position of the 'ON' and 'OFF' pixels relative to the edge of the ring electrode as a function of time. 'ON' pixels are defined here as pixels with an intensity over 110 au. Straight line fit between t = 1.6 s and 8.2 s to calculate the distance of change over time.

### REFERENCES

[1] Lerner, E.; Cordes, T.; Ingargiola, A.; Alhadid, Y.; Chung, S.; Michalet, X.; Weiss, S. Toward Dynamic Structural Biology: Two Decades of Single-Molecule Förster Resonance Energy Transfer. Science 2018, 359 (6373), eaan1133.

[2] Ambrose, B.; Baxter, J. M.; Cully, J.; et al. The smfBox Is an Open-Source Platform for Single-Molecule FRET. Nat. Commun. 2020, 11, 5641.
